# Deploying and *in vitro* gut model to assay the impact of a mannan-oligosaccharide prebiotic, Bio-MOS® on the Atlantic salmon (Salmo salar) gut microbiome

**DOI:** 10.1101/2020.10.07.328427

**Authors:** R. Kazlauskaite, B. Cheaib, J. Humble, C. Heys, U. Ijaz, S. Connelly, W.T. Sloan, J. Russell, L. Martinez-Rubio, J. Sweetman, A. Kitts, P. McGinnity, P. Lyons, M. Llewellyn

## Abstract

Mannose-oligosaccharide (MOS) pre-biotics are widely deployed in animal agriculture as immunomodulators as well as to enhance growth and gut health. Their mode of action is thought to be mediated through their impact on host microbial communities and associated metabolism. Bio-MOS is a commercially available pre-biotic currently used in the agri-feed industry. To assay Bio-MOS for potential use as a pre-biotic growth promotor in salmonid aquaculture, we modified an established Atlantic salmon *in vitro* gut model, SalmoSim, to evaluate its impact on host microbial communities. In biological triplicate, microbial communities were stabilised in SalmoSim followed by twenty-day exposure to the pre-biotic and then an eight day ‘wash out’ period. Exposure the MOS resulted in a significant increase in formate (p=0.001), propionate (p=0.037) and isovalerate (p=0.024) levels, correlated with increased abundances of several principally anaerobic microbial genera (*Fusobacteria, Agarivorans, Pseudoalteromonas, Myroides*). 16S rDNA sequencing confirmed a significant shift in microbial community composition in response to supplementation. In conjunction with previous *in vivo* studies linking enhanced VFA production alongside MOS supplementation to host growth and performance, our data suggest that Bio-MOS may be of value in salmonid production. Furthermore, our data highlight the potential role of *in vitro* gut models to augment *in vivo* trials of microbiome modulators.

## Introduction

Since the late 1970s, the salmon aquaculture sector has grown significantly, currently exceeding 1 million tonnes of salmon produced per year (FAO, 2018). Fish in aquaculture environments are exposed to events and interactions that are extensively different from the wild, such as changes in temperature and salinity, and close contact between animals that can favour potential disease outbreaks by proliferating pathogenic agents present from the surrounding environment from one animal to the other (Kennedy et al., 2016), as well as long term stress through aggression and overcrowding (Adams et al., 2007; Turnbull et al., 2005). Thus, the rapid expansion of the aquaculture sector requires tools that promote feed performance, reduce the need for medical treatments and reduce waste discharges while improving farmed fish quality and in this way boosting profitability.

In order to prevent disease outbreaks and improve feed performance, prebiotics are widely deployed in agriculture and aquaculture (Markowiak & Ślizewska, 2018; Patterson & Burkholder, 2003; Ringø et al., 2010). Prebiotics are defined as non-digestible food additives that have a beneficial effect on the host by stimulating growth and activity of the believed beneficial bacterial communities within gut thus improving overall animal health (Gibson, Glenn R. and Roberfroid, 1995). One of the most widely used and evaluated prebiotic types in aquaculture are mannooligosaccharides (MOS) that are made of glucomannoprotein-complexes derived from the outer layer of yeast cell walls (*Saccharomyces cerevisiae*) (Merrifield et al., 2010). MOS compounds were shown to improve gut function and health by increasing villi height, evenness and integrity in chickens (Hooge, 2004; Iji et al., 2001), cattle (Castillo et al., 2008) and fish (Dimitroglou et al., 2009). MOS supplementation in monogastrics (Halas & Nochta, 2012; Sims et al., 2004) has been reported to drive changes in host associated microbial communities. Associated increased volatile fatty acid production is reported and can have beneficial knock on effects in terms of host metabolism and gut health (Besten et al., 2013).

There are limited number of studies investigating the effect of MOS on the fish microbiome with disparities in the observed results that could be partially explained by the time of MOS supplementation, fish species, age or environmental conditions. For instance, it was found that MOS supplemented diets improved growth and/or feed utilization in some studies (Buentello et al., 2010; Gültepe et al., 2011; Staykov et al., 2007; Torrecillas et al., 2013; Yilmaz et al., 2007), but others found that MOS supplementation did not affect fish performance or feed efficiency (Peterson et al., 2010; Pryor et al., 2003; Razeghi Mansour et al., 2012). Detailed studies are needed to investigate the effect of MOS supplements on the fish microbiome to enhance our understand of the link between MOS and gut health. *In vitro* gut models offer the advantage of doing so in a replicated and controlled environment.

SalmoSim is a continuous salmon gut fermentation system aimed at simulating microbial communities present in the intestine of marine phase Atlantic Salmon (*Salmo salar*) (Kazlauskaite et al., 2020). The current study deploys a modified version of SalmoSim aims to evaluate the effect of Bio-MOS (Alltech), commercially available MOS product, on the microbial communities of the Atlantic salmon small intestine (pyloric caecum) in biological triplicate. We assayed microbial composition and fermentation in the SalmoSim system, showing a significant impact of Bio-MOS supplementation on both.

## Materials and Methods

### *In vivo* sample collection and *in vitro* system inoculation

Three adult Atlantic salmon (*Salmo salar*) gut samples were collected from the MOWI processing plant in Fort William, and transfer in the anaerobic box on ice. Once samples were brought back to the laboratory, they were placed in the anaerobic hood and gut contents from pyloric caeca compartment were scraped and collected to the separate tubes. Half of the content of these tubes was stored in −80°C freezer, while other half was used as an inoculum for SalmoSim system. Fresh bacterial inoculums were prepared for the *in vitro* trial from the pyloric caeca compartment sampled from individual fish (three biological replicates). Prior to inoculation, inoculums were dissolved in 1 ml of autoclaved 35 g/L Instant Ocean^®^ Sea Salt solution.

### SalmoSim *in vitro* system preparation

*In vitro* system feed media was prepared by combining the following for a total of 2 litres: 35 g/L of Instant Ocean^®^ Sea Salt, 10 g/L of the Fish meal, 1 g/L freeze-dried mucous collected from the pyloric caecum, 2 litres of deionised water and 0.4% of Bio-MOS for the prebiotic supplemented feed. This feed was then autoclave-sterilised, followed by sieving of the bulky flocculate, and finally subjected to the second round of autoclaving. System architecture was prepared as described previously with some modifications (Kazlauskaite et al., 2020).In short, appropriate tubes and probes were attached to a one 2 litre double-jacketed bioreactor and three 500 ml Applikon Mini Bioreactors filled with four 1cm^3^ cubes made from aquarium sponge filters, autoclave sterilised and connected as in Figure 1A. Nitrogen gas was periodically bubbled through each vessel to maintain anaerobic conditions. The two litre double jacketed bioreactor and three 500 ml bioreactors were filled with 1.5 litres and 400 ml of feed media respectively. Once the system was set up, media transfer, gas flow and acid/base addition were undertaken for twenty-four hours axenically in order to stabilise the temperature, pH, and oxygen concentration with respect to levels measured from adult salmon. SalmoSim system diagram is visualised in Figure 1 A. Physiochemical conditions within three 500 ml SalmoSim gut compartments were kept similar to the values measured *in vivo* (Kazlauskaite et al., 2020): temperature inside the reactor vessels was maintained at 12°C, dissolved oxygen contents were kept at 0% by daily flushing with N2 gas for 20 minutes, and pH 7.0 by the addition of 0.01M NaOH and 0.01M HCl. The 2 litre double jacketed bioreactor (represents sterile stomach compartment) was kept at 12°C and pH at 4.0 by the addition of 0.01M HCl. During this experiment (apart from the initial pre-growth period) transfer rate of slurry between reactor vessels was 238 ml per day. Finally, 1 ml of filtered salmon bile and 0.5 ml of autoclaved 5% mucous solution were added to the three bioreactors simulating pyloric caecum compartments on a daily basis.

**Figure 1.**
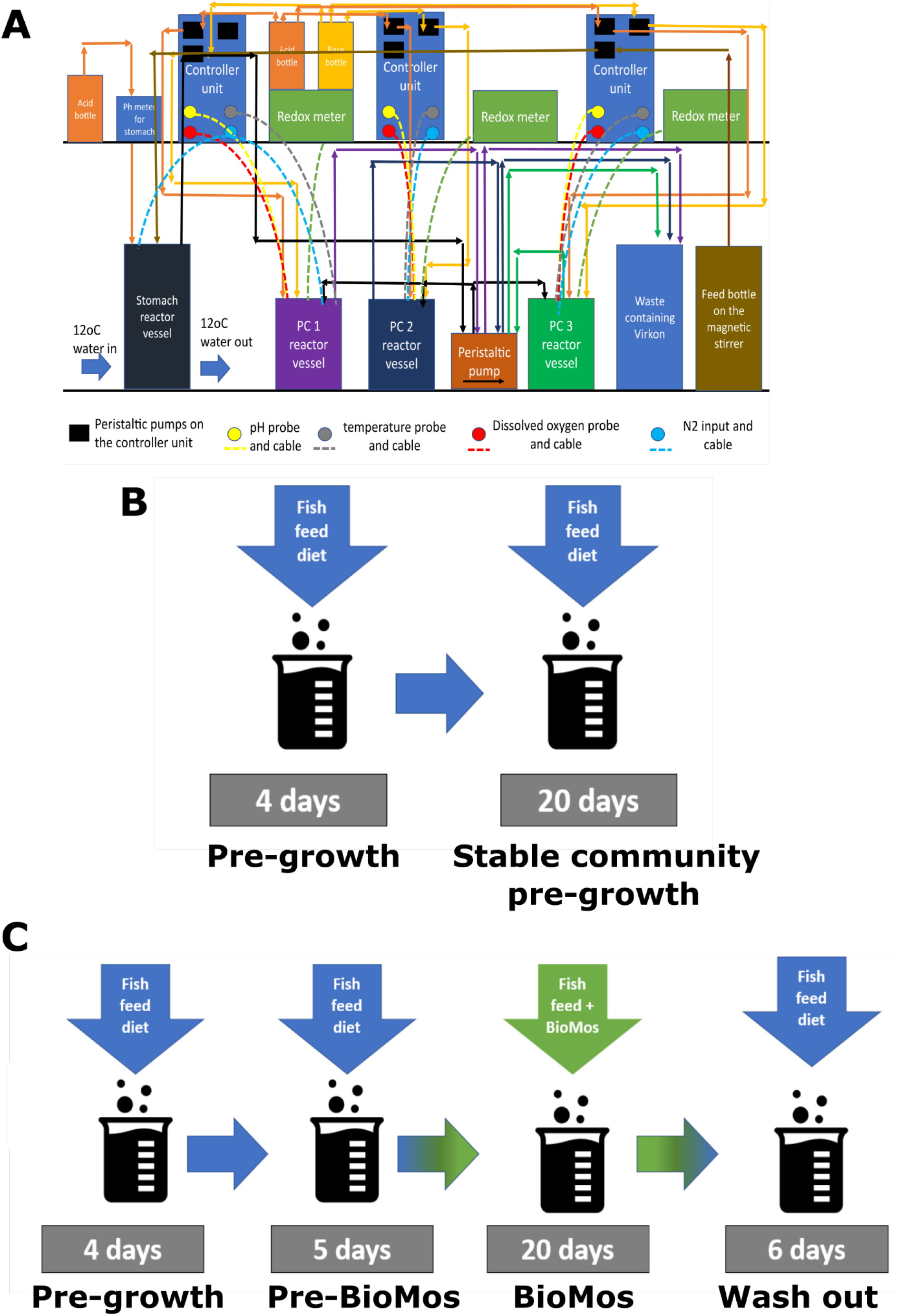
Artificial gut model system set-up and in vitro trial set up. Figure 1 Summarises the experimental study design. 1A is a schematic representation of SalmoSim system; 1B visual representation of stable community pre-growth run within SalmoSim system; 1C visual representation of the main experimental run that involved four stages: (i) pre-growth (without feed transfer for 4 days), (ii) feeding system with Fish meal (Pre-BioMOS: 5 days), (iii) feeding system with Fish meal diet supplemented with Bio-MOS (BioMOS: 20 days), (iv) wash out period during which system was fed Fish meal without the addition of prebiotic (Wash out: 6 days).

### SalmoSim inoculation and microbial growth

To generate stable and representative microbial communities for experimentation (Kazlauskaite et al., 2020), microbial communities were grown for twenty-four days in advanced of the experiment As such, fresh inocula from pyloric caeca were added to three 500 ml bioreactors and pre-grown for 4 days without media transfer, followed by 20 days feeding the system at a 238 ml per day feed transfer rate. Fifteen ml of the stable communities were collected at the end of this pre-growth period, centrifuged at 3000g for 10 minutes and supernatant removed. The pellet was then dissolved in 1 ml of autoclaved 35 g/L Instant Ocean^®^ Sea Salt solution, flash frozen in liquid nitrogen for 5 minutes and stored long term in −80°C freezer.

### Assaying Bio-MOS impact on microbial communities in the SalmoSim *in vitro* system

Frozen pre-grown stable pyloric caeca samples were thrawn on ice and added to the SalmoSim system (into each 500 ml bioreactor bacterial communities from a different fish were added). The system was run in several stages: (i) 4-day initial pre-growth period without feed transfer (Pre-growth), (ii) 5-day period during which SalmoSim was fed on feed without prebiotic (Pre-BioMOS), (iii) 20-day period during which SalmoSim was fed on feed with BioMos (BioMOS), (iv) 6-day wash out period during which SalmoSim was fed on Fish meal diet without addition of prebiotic (Wash out). The schematical representation of the main experimental design is visually represented in Figure 1 C. Sixteeen samples were collected throughout the experimental run as described previously (Kazlauskaite et al., 2020).

### Genomic DNA extraction and NGS library preparation

The DNA extraction and NGS library preparation protocol was previously described (Heys et al., 2020; Kazlauskaite et al., 2020). In short, the samples collected from SalmoSim system and stable pre-grown inoculums were thrawn on ice and exposed to bead-beating step for 60 seconds by combining samples with MP Biomedicals™ 1/4” CERAMIC SPHERE (Thermo Fisher Scientific, USA) and Lysing Matrix A Bulk (MP Biomedicals, USA). Later, DNA was extracted by using the QIAamp^®^ DNA Stool kit (Qiagen, Valencia, CA, USA) according to the manufacturer’s protocol (Claassen et al., 2013). After, extracted DNA was amplified by using primers targeting V1 bacterial rDNA 16s region under the following PCR conditions: 95°C for ten minutes, followed by 25 cycles at 95°C for 30 seconds, 55°C for 30 seconds and 72°C for 30 seconds, followed by a final elongation step of 72°C for 10 minutes. The second-round PCR, which enabled the addition of the external multiplex identifiers (barcodes), involved only six cycles and otherwise identical reaction conditions to the first round PCR. This was followed by the DNA clean-up by using Agencourt AMPure XP beads (Beckman Coulter, USA) according to the manufacturers’ protocol and gel-purification using the QIAquick Gel Extraction Kit (Qiagen, Valencia, CA, USA). All the PCR products were pulled together at 10nM concentration and sent for the HiSeq 2500 sequencing.

### NGS data analysis

NGS data was analysed as described previously (Kazlauskaite et al., 2020). In short, two alpha diversity metrics were calculated effective microbial richness and effective Shannon diversity by using Rhea pipeline (Lagkouvardos et al., 2017b) and visualised by using microbiomeSeq package based on phyloseq package (McMurdie & Holmes, 2013; Ssekagiri et al., 2017).

Differential abundance was calculated by using microbiomeSeq package based on DESeq2 package (Love et al., 2017; Ssekagiri et al., 2017). BIOM generated OTU table was used as an input to calculate differentially abundant OTUs between different experimental phases (Pre-BioMOS, BioMOS and Wash out) based on the Negative Binomial (Gamma-Poisson) distribution.

Pearson correlation coefficient was calculated between taxonomic variables (OTUs) and two different datasets of meta-variables: (i) ammonia and protein concentrations and (ii) measured VFA values. All these correlations were calculated and visualised by using tools supplied by Rhea pipeline (Lagkouvardos et al., 2017a).

Principle Coordinates Analysis (PCoA) was performed by using microbiomeSeq package based on phyloseq package (Love et al., 2017; Ssekagiri et al., 2017) with Bray-Curtis dissimilarity measures calculated by using the vegdist() function from the vegan v2.4-2 package (Oksanen et al., 2013). Bray-Curtis distances were calculated for four different datasets: the full dataset (containing all biological replicates together), and, three different subsets each containing only one of the three biological replicate samples from SalmoSim: Fish 1, 2, or 3.

Finally, beta diversity was calculated for two different datasets: all (completed data set containing all the samples sequenced) and subset (containing all samples for Pre-BioMOS and Wash out period, but only stable samplings from BioMOS period (time points 11, 12 and 13)). These datasets were then used to compute ecological distances by using Bray-Curtis and Jaccards method for each subset shown in table 3 by using vegdist() function from the vegan v2.4-2 package (Oksanen et al., 2013). Furthermore, the phylogenetical distances were computed for each dataset by using GUniFrac() function (generalised UniFrac) from the Rhea package at the 0% (unweighted), 50% (balanced) and 100% (weighted) weight on the abundant lineages within the phylogenetic tree (Lagkouvardos et al., 2017a). These both ecological and phylogenetical distances were then visualised in two dimensions by Multi-Dimensional Scaling (MDS) and non-metric MDS (NMDS) (Anderson, 2001). Finally, a permutational multivariate analysis of variance (PERMANOVA) by using calculated ecological and phylogenetic distances was performed to determine if the separation of selected groups is significant as a whole and in pairs (Anderson, 2001).

### Protein fermentation and Volatile Fatty Acid (VFA) analysis

At each sampling point, microbial protein fermentation was assessed by measuring the protein concentration using Thermo Scientific™ Pierce™ BCA Protein Assay Kit (Thermo Fisher Scientific, USA) and the ammonia concentration using Sigma-Aldrich^®^ Ammonia Assay Kit (Sigma-Aldrich, USA). Both methods were performed according to manufacturer protocol by using a Jenway 6305 UV/Visible Spectrophotometer (Jenway, USA). For VFA analysis, nine samples from each pyloric caecum compartment were collected (from 3 biological replicates): 3 samples from Pre-BioMOS period (days 2-6), 3 samples from stable time points from the period while SalmoSim was fed on feed supplemented with Bio-MOS (days 22-26), and 3 samples from the Wash out period (days 28-32). VFA sampling was performed as described previously (Kazlauskaite et al., 2020). Extracted VFAs were sent for gas chromatographic analysis at the MS-Omics (Denmark).

In order to establish whether VFA concentrations are statistically different between different experimental phases (Pre-BioMOS, BioMOS and Wash out), a linear mixed effect model was deployed (Model 1) treating time point (sampling time point) and run (biological replicate of SalmoSim system) as random effects.

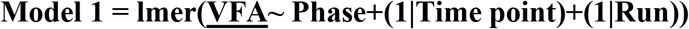

Finally, in order to establish whether ammonia production statistically changes throughout experimental run, a linear mixed effect model was deployed (Model 2) treating run (biological replicate of SalmoSim system) as random effect.

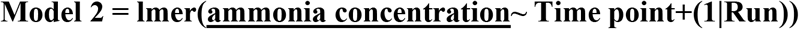

## Results

In order to explore the impact of the MOS prebiotic on microbial communities in SalmoSim, microbes in different experimental phases (Pre-BioMOS, BioMOS and wash out) were surveyed using Illumina NovaSeq amplicon sequencing of the 16S V1 rDNA locus. In total 11.5 million sequence reads were obtained after quality filtering. Alpha diversity metrics (Effective richness (Figure 2 A) and effective Shannon diversity (Figure 2 B)) indicated that the initial inoculum contained the lowest number of OTUs and had the lowest bacterial richness compared to later sampling time points from SalmoSim system, but these differences were not statistically significant. Furthermore, this figure indicates no statistically significant differences between different experimental phases in both terms of effective richness and Shannon diversity. Taken together, diversity and richness estimates suggest non-statistically significant increase of microbial taxa as a result of transfer into SalmoSim system, but overwise stable diversity and richness over the different experimental phases.

**Figure 2.**
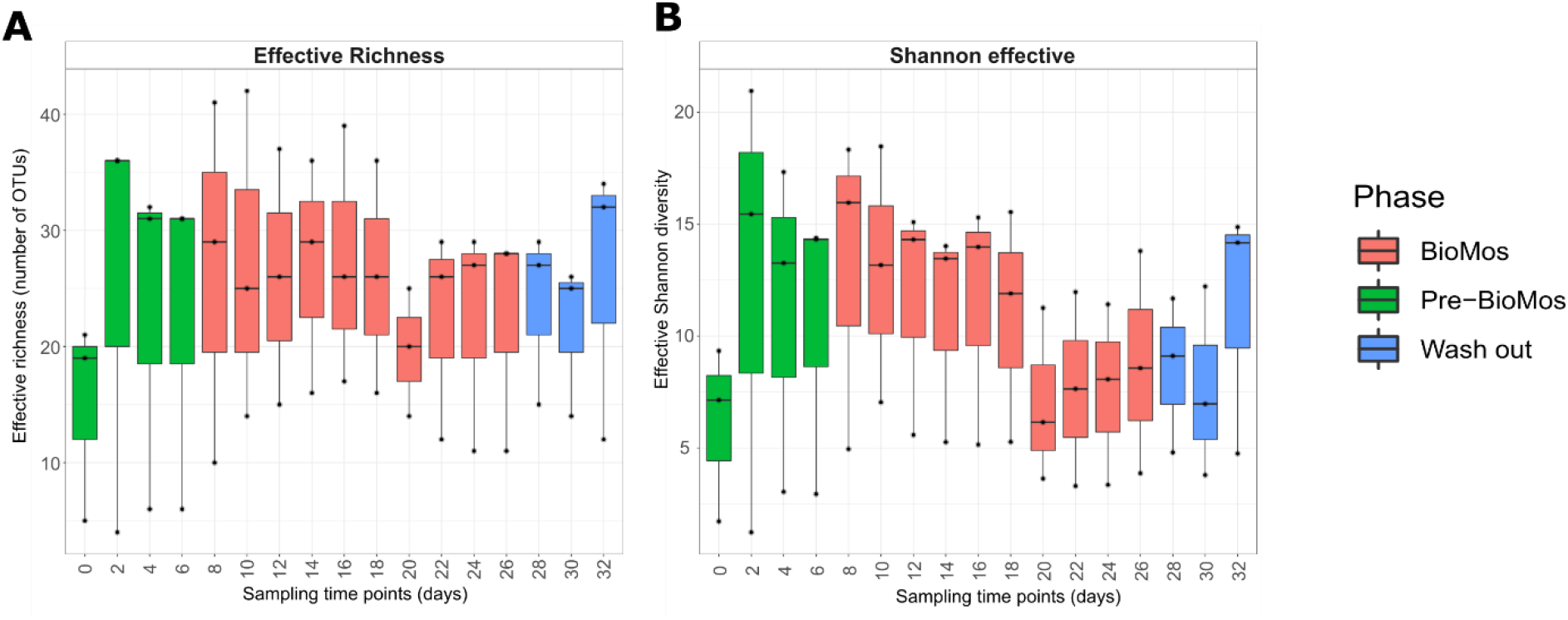
Alpha-diversity dynamics within the SalmoSim system during exposure to Bio-MOS prebiotics. The figure represents different alpha diversity outputs at different sampling time points (days) from SalmoSim system. Time point 0 represents microbial community composition within initial SalmoSim inoculum from the pre-grown stable bacterial communities, time points 2-6 identifies samples from SalmoSim system fed on Fish meal diet alone (Pre-BioMOS: green), time points 8-26 identifies samples from SalmoSim system fed on Fish meal diet with addition of Bio-MOS (BioMOS: red), and time points 28-32 identifies samples from wash out period while SalmoSim was fed on feed without addition of prebiotic (Wash out: blue). **A** visually represents effective richness (number of OTUs), **B** represents effective Shannon diversity.

To provide an overview of microbial composition and variation in the experiment, a PCoA plot was constructed based on Bray-Curtis distanced between samples (Figure 3 A-D). Biological replicate (the founding inoculum of each SalmoSim run) appears to be a major driver of community composition in the experiment (Figure 3 A). This is supported by Figure 4 that visually represents varying microbial composition within different fish. Only once individual SalmoSim replicates are visualised separately do the changes to microbial communities in response to the different experimental conditions become apparent (Figures 3 B-D) and reflect PERMANOVAs carried out in Table 1. These results indicate that bacterial communities shift from Pre-BioMOS to BioMOS, but they remain fairly stable (statistically similar) between BioMOS and Wash out periods, however, not necessarily along the same axes as each SalmoSim sample indicative, perhaps, or a different microbiological basis for that change. This trend is confirmed in Figure 5 that indicates a more substantial shift in microbial community profile between Pre-BioMOS and BioMOS phases in Fish 2 and 3, but to the lesser extent in Fish 1. These results were confirmed by performing beta-diversity analysis using both phylogenetic and ecological distances, both of which indicated statistically significant differences between Pre-BioMOS and BioMOS phases, but not between BioMOS and Wash out periods (Table 1). To investigate this further differential abundance analysis was performed, which indicated that only 1 OTU belonging to *Psychrobacter* genus was significantly differentially abundant between the Pre-BioMOS and BioMOS phases across all biological replicates (Figure 5). Furthermore, 1 OTU also belonging to *Psychrobacter* genus was differentially abundant between BioMos and Wash out phases (Figure 5).

**Figure 3.**
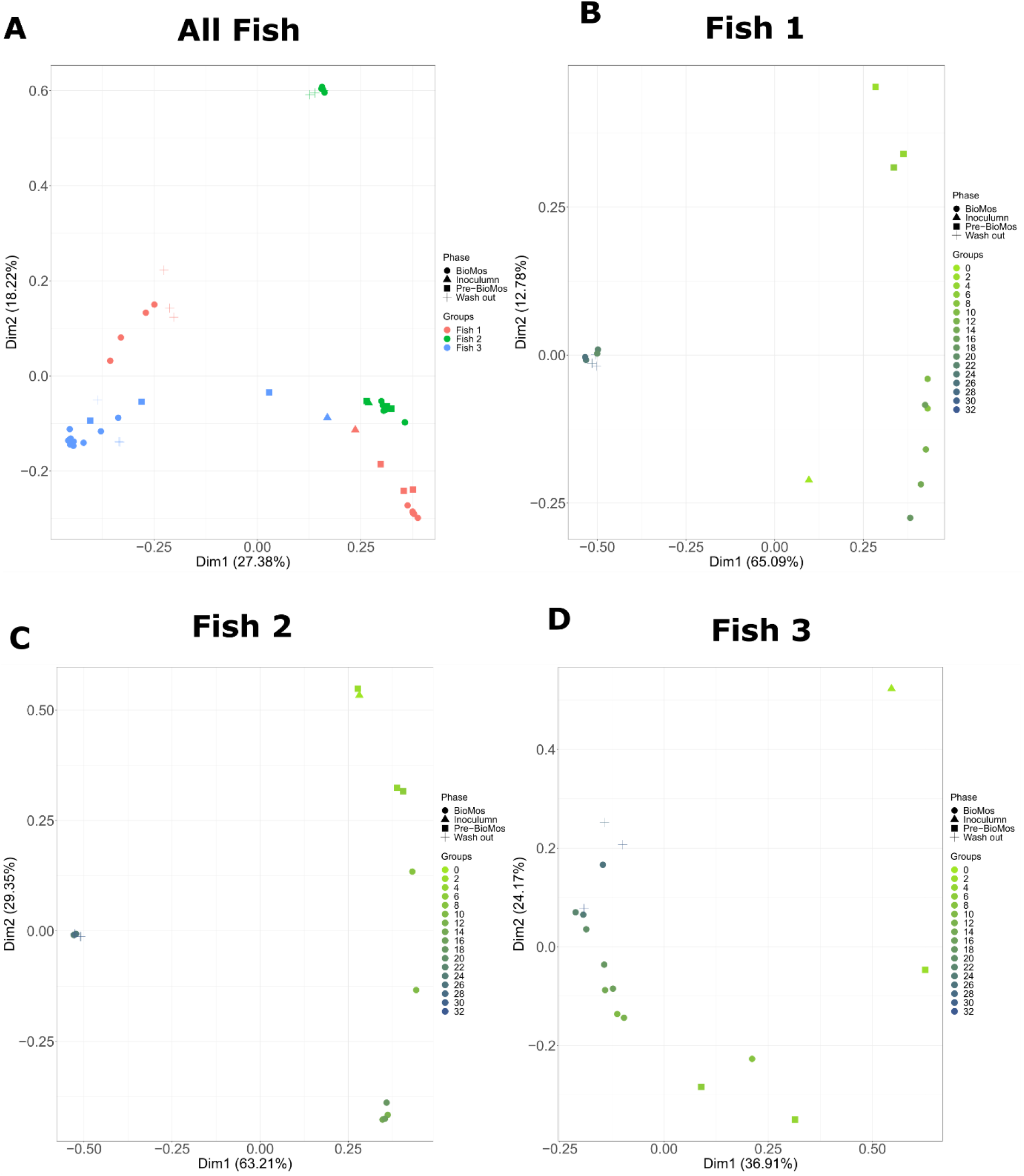
PCoA plots visualising bacterial communities within SalmoSim bioreactors during exposure to Bio-MOS prebiotic. Figure above visualises four principal-coordinate analysis (PCoA) plots for Bray-Curtis dissimilarity measures for different experimental phases (Inoculum, Pre-BioMOS, BioMOS and Wash out), different sampling time points from SalmoSim system and different biological replicates. **A** represents all sequenced data together for all 3 biological replicates in which different colours represent different biological replicates (samples from pyloric ceacum from 3 different fish) and different shapes represent different experimental phases(Inoculum, Pre-BioMOS, BioMOS and Wash out); **B-D** represent sequenced data for each individual biological replicate (**B**: Fish 1, **C**: Fish 2, **D**: Fish 3). In figures B-D different colours represent different sampling time points and different shapes represent different experimental phases (Inoculum, Pre-BioMOS, BioMOS and Wash out). Dim 1 is principal coordinate 1, and Dim 2 is principle coordinate 2.

**Figure 4.**
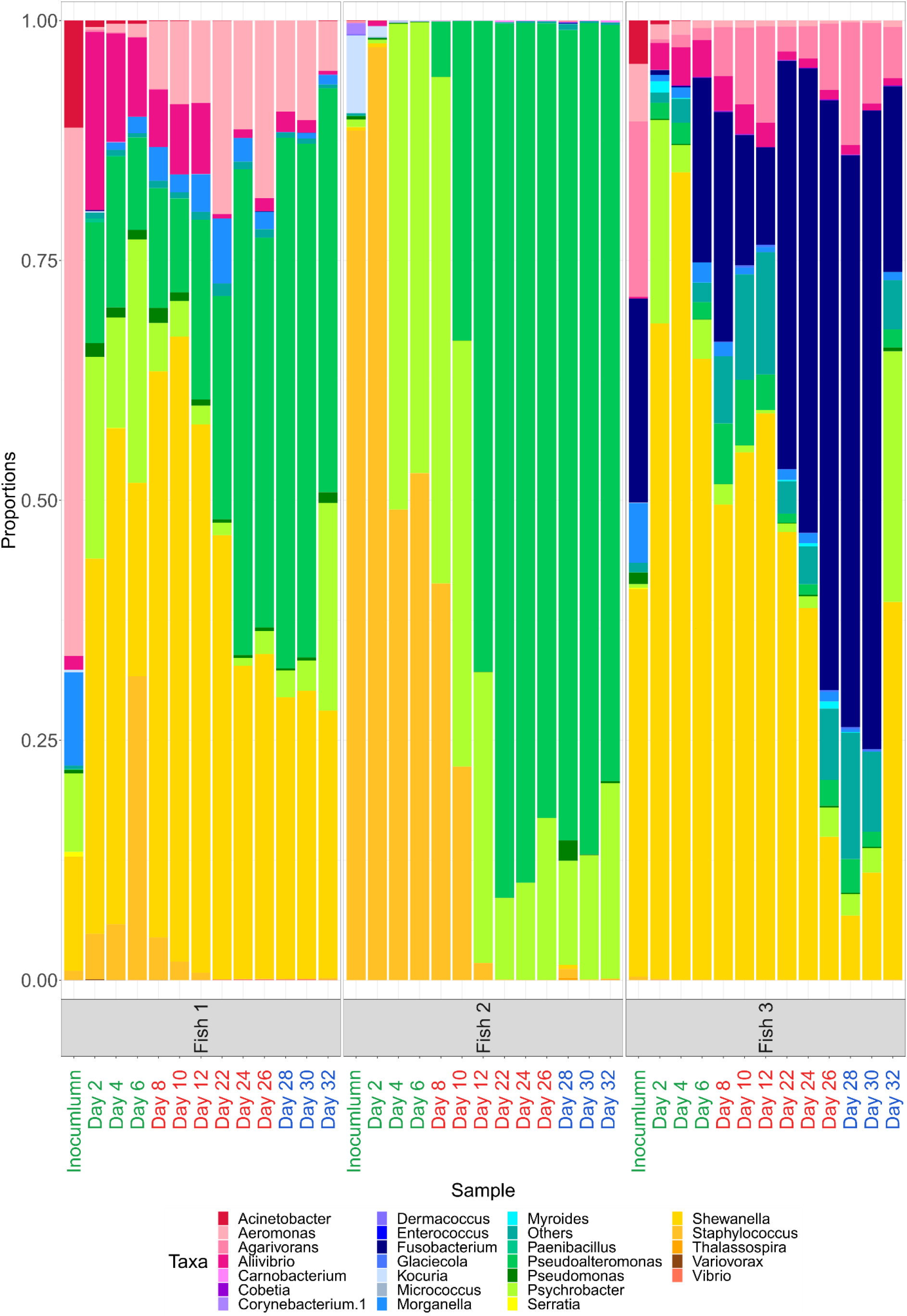
Microbial composition (25 most common genus + others) amongst different biological replicates and experimental phases. Labels on X axis in green represent samples from Pre-BioMOS phases, in red samples fed on BioMOS phase and in blue samples from Wash out period. Only subset of time points is visualised for each phase: time points 2-6 for Pre-BioMOS, 8-12 and 22-24 for BioMOS, and 28-32 for Wash out.

**Figure 5.**
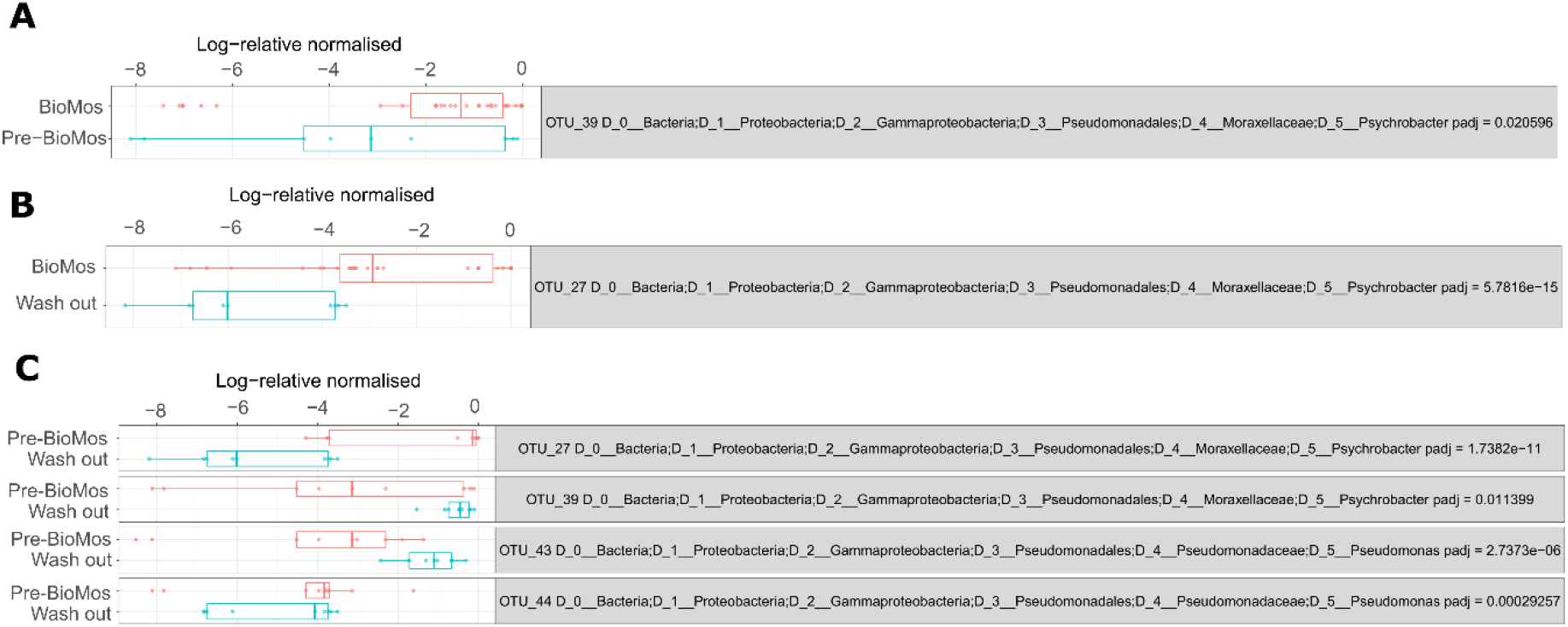
Differential abundance of OTUs between different experimental phases (Pre-BioMOS, BioMOS and Wash out) Differential abundant OTUs between different experimental phases: Pre-BioMOS vs BioMOS (Figure 5 A), Bio-MOS vs Wash out (Figure 5 B), Pre-BioMOS vs Wash out (Figure 5 C).

**Table 1.** Beta diversity and differential abundance values from the comparison of microbial composition between different phases (Pre-BioMOS, BioMOS and Wash out) The table summarises different beta-diversity analysis outputs calculated by using different distances: phylogenetic (unweighted, balanced and weighted UniFrac) and ecological (Bray-Curtis and Jaccard’s), between different experimental phases: Pre-BioMOS, BioMOS and Wash out. Numbers represent p-values, with p-values <0.05 identifying statistically significant differences between compared groups. The comparisons are shown for 3 different datasets: All (completed data set containing all the samples sequenced) and Subset (containing all samples for Pre-BioMOS and Wash out period, but only stable samplings from BioMOS period (time points 11, 12 and 13)).

| Test |  | Data | Pre-BioMOS vs BioMOS | BioMOS vs Wash out | Pre-BioMOS vs Wash out |
| --- | --- | --- | --- | --- | --- |
| UniFrac | Unweighted (0%) | All | 0.042 | 0.029 | 0.001 |
|  |  | Subset | 0.002 | 0.184 | 0.001 |
|  | Balanced (50%) | All | 0.023 | 0.207 | 0.001 |
|  |  | Subset | 0.007 | 0.648 | 0.001 |
|  | Weighted (100%) | All | 0.022 | 0.37 | 0.002 |
|  |  | Subset | 0.002 | 0.717 | 0.001 |
| Bray-Curtis |  | All | 0.034 | 0.04 | 0.001 |
|  |  | Subset | 0.003 | 0.727 | 0.001 |
| Jaccards |  | All | 0.018 | 0.05 | 0.001 |
|  |  | Subset | 0.001 | 0.8 | 0.001 |
| Differential abundance at OTU level |  | Number of different OTUs between comparisons | 1 | 1 | 4 |

VFA levels were measured throughout the SalmoSim trial for the stable time points (time points 2, 6 and 8 for Pre-BioMOS, time points 22, 24 and 26 for BioMOS, and time points 28, 30 and 32 for Wash out period). These results are visually represented in Figure 6 and indicate that statistically significant increases were found between Pre-BioMOS and BioMOS phases in formic, propanoic and 3-methylbutanoic acid concentrations. No significant differences in any VFA production by the system was noted between BioMOS and Wash out periods. To assay the potential microbial drivers of, or microbial correlates with, VFA production, pearson correlation coefficients were calculated between these and individual OTU abundances. Data are shown in Figure 7 where multiple instanced on strong (R>0.8) positive and negative correlations were noted with individual OTUs. Acetate and propionate showed the strongest individual-level correlations. Finally, Figure 8 visually summarises measured ammonia (NH_3_) concentration changes through experiment. The data indicates statistically significant increase in ammonia production between time points 2 and 4, and between time points 20 and 22, and decrease in ammonia concentration between time points 30 and 32.

**Figure 6.**
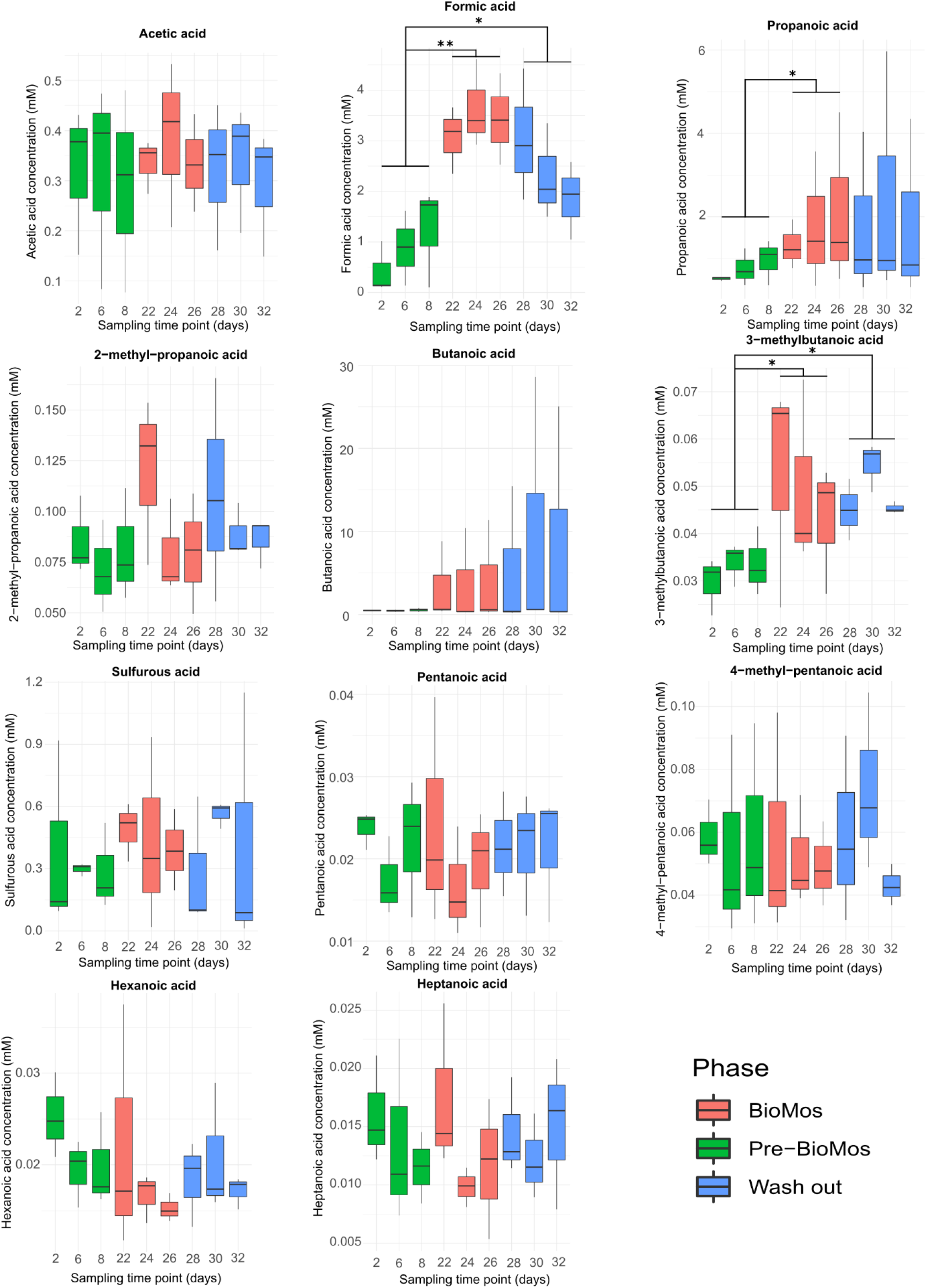
VFA responses in SalmoSim pyloric caecum compartment after Bio-MOS introduction and subsequent wash out period. Figure above visually represents 11 volatile fatty acid production in three different experimental phases: (i) SalmoSim fed on Fish meal alone without prebiotic addition (Pre-BioMOS: green), (ii) SalmoSim fed on Fish meal with addition of Bio-MOS (BioMOS: red), (iii) wash out period during which SalmoSim was fed on Fish meal without Bio-MOS (Wash out: blue). X axis represents the concentration of specific volatile fatty acid (mM) while the Y axis represents different sampling time points (days). The lines above bar plots represent statistically significant differences between different experimental phases. The stars flag the levels of significance: one star (*) for p-values between 0.05 and 0.01, two stars (**) for p-values between 0.01 and 0.001, and three stars (***) for p-values below 0.001.

**Figure 7.**
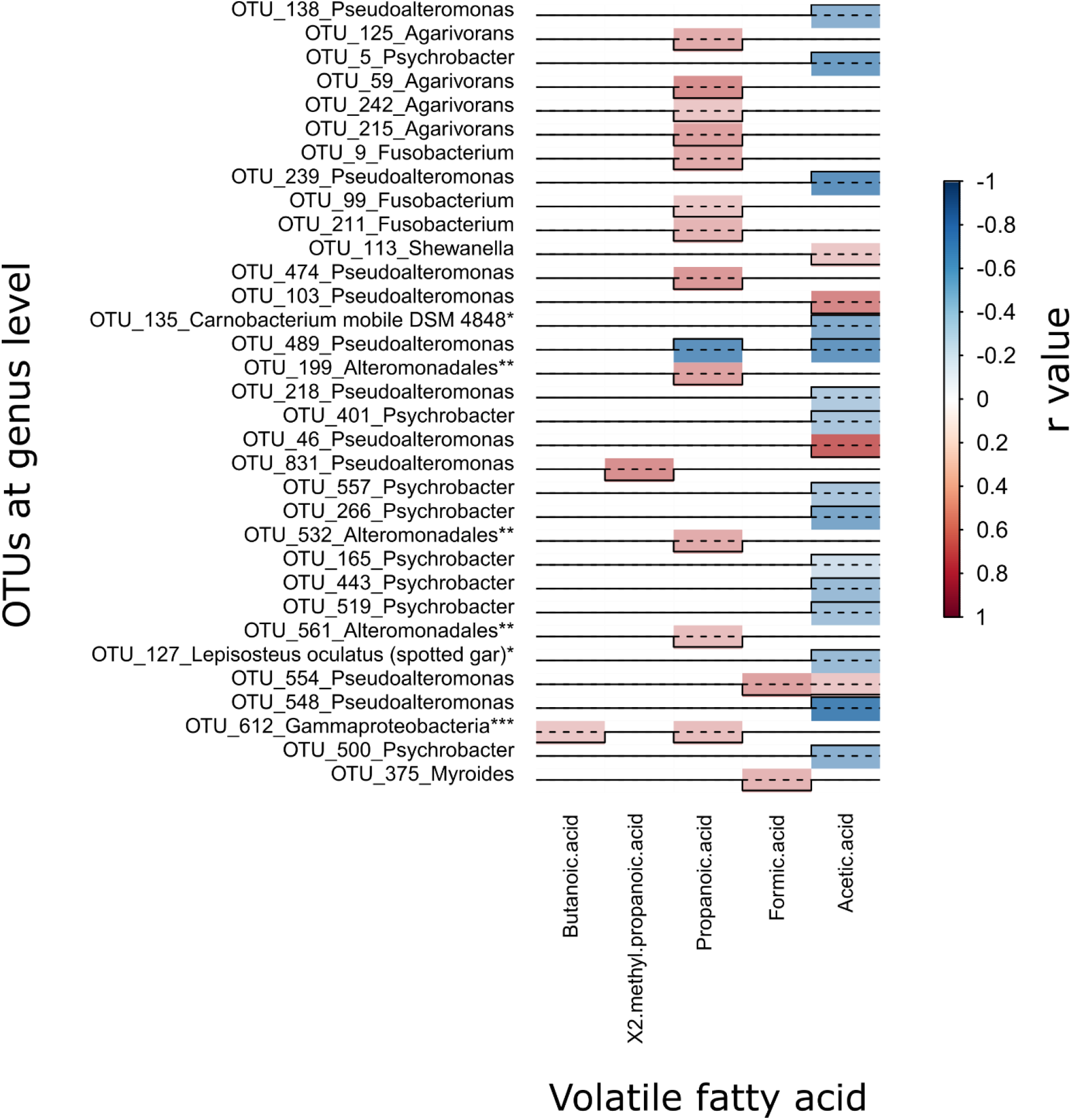
Person correlation coefficients across different values measured and taxonomic variables. Figure visually summarises calculated statistically significant (p<0.05) and strongly correlated (r>0.8) Pearson correlation coefficients across a set of meta- and taxonomic variables. Blue colour represents negative correlation and red colour represents positive correlations, with 1 being strong positive correlation and −1 being strong negative correlation. All OTUs are summarised to genus level apart from * to species level, ** to order and *** to class level.

**Figure 8.**
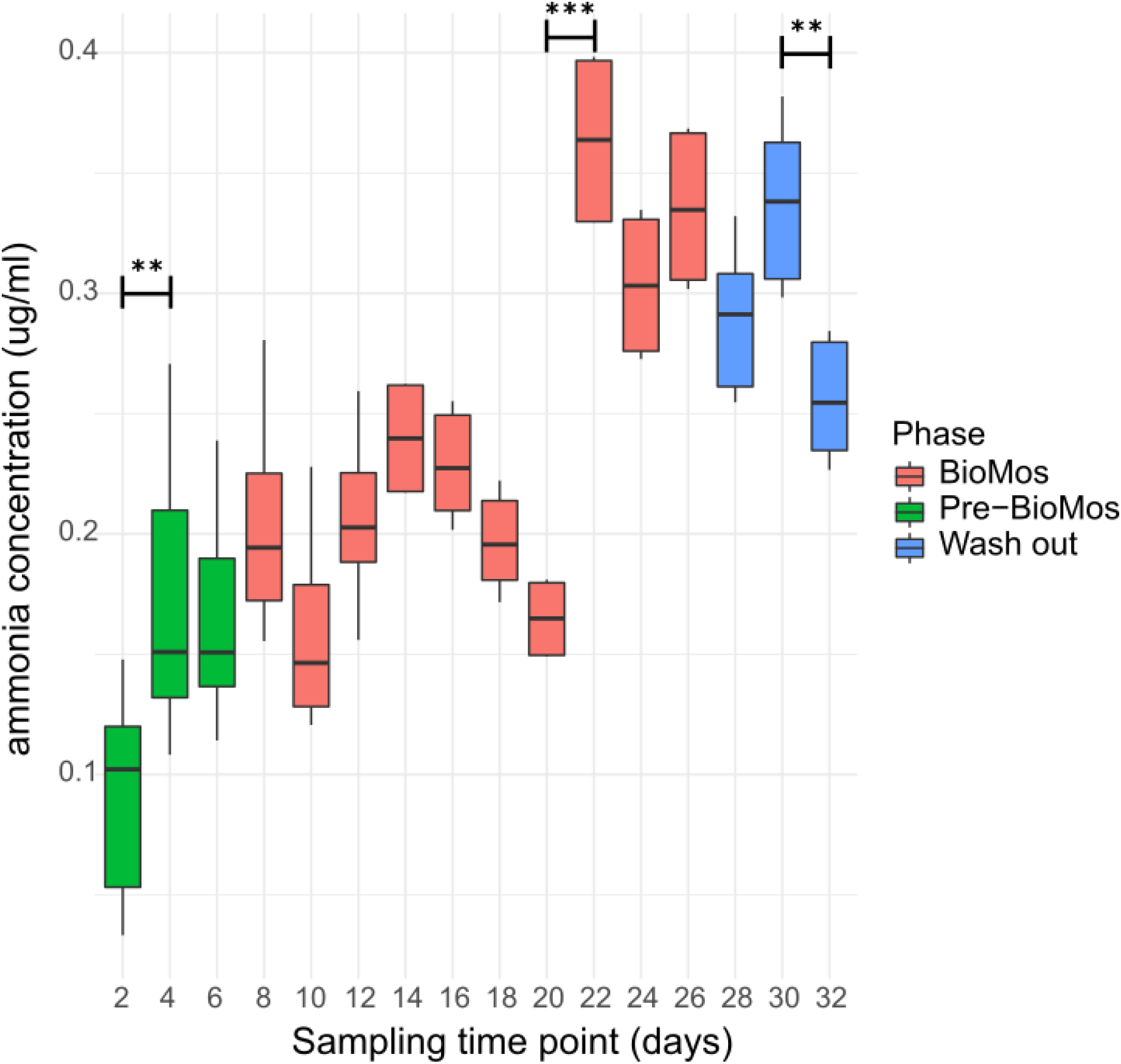
ammonia (NH_3_) concentration in SalmoSim pyloric caecum compartment throughout experiment. Figure above visually represents ammonia (NH_3_) production in three different experimental phases: (i) SalmoSim fed on Fish meal alone without prebiotic addition (Pre-BioMOS: green), (ii) SalmoSim fed on Fish meal with addition of Bio-MOS (BioMOS: red), (iii) wash out period during which SalmoSim was fed on Fish meal without Bio-MOS (Wash out: blue). X axis represents the concentration of ammonia (μg/ml) while the Y axis represents different sampling time points (days). The lines above bar plots represent statistically significant differences between sequential time points. The stars flag the levels of significance: one star (*) for p-values between 0.05 and 0.01, two stars (**) for p-values between 0.01 and 0.001, and three stars (***) for p-values below 0.001.

## Discussion

Our study aimed to evaluate the effect of commercially available MOS product (Bio-MOS) on the microbial communities within Atlantic salmon gut using newly developed artificial salmon gut simulator SalmoSim. Inclusion of Bio-MOS in feed did not affect microbial community diversity and richness in the SalmoSim system, nor did subsequent removal of the prebiotic during wash out.). Biological replicate (the founding inoculum of each SalmoSim run) appears to be a major driver of community composition in the experiment. Our results indicate that bacterial communities diverged in terms of their composition between Pre-Bio-MOS and Bio-MOS experimental phases but were statistically similar between Bio-MOS and wash out periods. Similar trends were identified in bacterial activity (VFA production) that showed statistically significant increases in formic, propanoic and 3-methylbutanoic acid concentrations during the shift from Pre-Bio-MOS and Bio-MOS phase, but no statistically significant change in bacterial activity between Bio-MOS and wash-out periods. Finally, a statistically significant increase in the ammonia production during BioMOS phase was observed at the later time points (between days 20 and 22), followed in the reduction in ammonia concentration during Wash out period (between days 30 and 32).

Several studies indicate that in vertebrates supplementing feed with MOS increases the production of propionate and butyrate by gut bacteria (Ao & Choct, 2013; Pan et al., 2009; Zdunczyk et al., 2005), while other studies find no effect of MOS on the VFA production (Gürbüz et al., 2010). In our study we saw a statistically significant increase in the production of formic, propanoic and 3-methylbutanoic acids in the SalmoSim system upon the switch to the feed supplemented with Bio-MOS. Propionate is commonly absorbed and metabolised by the liver, where it impacts host physiology via regulation of energy metabolism in humans (El Hage et al., 2020) and is associated with healthy gut histological development and enhanced growth in fish and shellfish (da Silva et al., 2016; Wassef et al., 2020). Formic acid, although frequently deployed as an acidifier in monogastrics to limit the growth of enteric pathogens (Luise et al., 2020), is not known to directly impact host phenotype. Similarly, isovaleric (3-methylbutanoic) acid is not known to directly impact host phenotype, except as the rare genetic disorder that occurs in humans, isovaleric acidemia, where the compound accumulates at high levels in the absence of isovaleric acid-CoA dehydrogenase activity in host tissues (Vockley & Ensenauer, 2006).

Alongside the effect of Bio-MOS, our results indicate strong and statistically significant positive correlation between the concentration of propanoic acid and OTUs belonging to multiple genera: Agarivorans (facultative anaerobic), Fusobacterium (anaerobic), Pseudoalteromonas (facultative anaerobic). Alongside the increase in formic acid as a result of Bio-MOS, OTUs belonging to Pseudoalteromonas (facultative anaerobic) and Myroides (aerobic) genus also increased in abundance. The causal directionality between these genera and the respective VFAs is hard to establish. Previously, a positive correlation was found between Fusobacterium genus and propanoic acid concentration in humans (Riordan, 2007). Propionate is a key substrate that can metabolised by several classes of methanogenic anaerobes (Mah et al., 1990) and may be driving the growth of the genera noted here. Equally, propionate is a major product of microbial metabolism of amino acids (Louis & Flint, 2017), and more efficient protein metabolism in the system by certain genera is likely drive its abundance. Ammoniacal nitrogen production was noted to increase after Bio-MOS has been added, albeit with a noticeable lag. Furthermore, although formate, propionate, isovalerate and ammonia show a downward trend after the remove of Bio-MOS, it is clear that a longer washout period is required to allow VFA and ammonia levels to recover their pre-Bio-MOS levels.

Research suggests that feed supplementation with MOS helps to improve the immune response in animals by stimulation of the production of mannose-binding proteins which are involved in phagocytosis and activation of the complement system (Franklin et al., 2005; Taschuk & Griebel, 2012). Such relationships with host immunity are difficult to predict with a simplified *in vitro* system. It is thought the feed supplementation with MOS elevates the immune response in host by increasing the lactic acid bacteria (LAB) levels in common carp (Momeni-Moghaddam et al., 2015). However, we did not observe any changes in LAB which could be explained by their low abundance in the initial inoculum. A fishmeal-based diet with limited carbohydrate content was used to perform this experiment, and has been previously linked to lower abundances of lactic acid producing bacteria when compared to microbial gut composition of Atlantic salmon fed on plant-based feed (Reveco et al., 2014). To enhance LAB growth alongside MOS in protein rich diets, some carbohydrate supplementation may be necessary.

Our study indicates the positive correlation between Bio-MOS supplementation and production of propanoic and formic acids, both of which are known to benefit animal microbiome and health (EFSA, 2014; Haque et al., 1970). Although, our *in vitro* model lacks host component, previous studies involving the use of gut simulators to analyse the effectiveness of various pre-biotics were shown to produce similar results to *in vivo* trials (Duysburgh et al., 2020; Sivieri et al., 2014). Thus, highlighting the usefulness of various in vitro gut systems, such as SalmoSim, to study the effectiveness of various feed additives.

